# How lucky was the genetic investigation in the Golden State Killer case?

**DOI:** 10.1101/531384

**Authors:** Michael (Doc) Edge, Graham Coop

**Affiliations:** Center for Population Biology, University of California, Davis; Department of Evolution and Ecology, University of California, Davis

## Abstract

Long-range forensic familial searching is a new method in forensic genetics. In long-range search, a sample of interest is genotyped at single-nucleotide polymorphism (SNP) markers, and the genotype is compared with a large database in order to find relatives. Here, we perform some simple calculations that explore the basic phenomena that govern long-range searching. Two opposing phenomena—one genealogical and one genetic—govern the success of the search in a database of a given size. As one considers more distant genealogical relationships, any target sample is likely to have more relatives—on average, one has more second cousins than first cousins, and so on. But more distant relatives are also harder to detect genetically. Starting with third cousins, there is an appreciable chance that a given genealogical relationship will not be detectable genetically. Given the balance of these genealogical and genetic phenomena and the size of databases currently queryable by law enforcement, it is likely that most people with substantial recent ancestry in the United States are accessible via long-range search.

**Note:** This material was originally posted on the Coop lab site on May 7th, 2018, soon after the reporting of the arrest of Joseph DeAngelo in the Golden State Killer case, one of the first high-profile uses of long-range familial search. Subsequently, Erlich et al. (2018) published a detailed analysis in a large empirical dataset along with a theoretical analysis of a model similar to the one we use here, obtaining results broadly consistent with the ones presented here. Because Erlich and colleagues kindly cited this work when describing their model, we thought it would be appropriate to post this material in a venue where it is more easily cited.

---

On April 24th, 2018, police arrested Joseph DeAngelo as a suspect in case of the Golden State Killer, an infamous serial murderer and rapist whose case has been open for over forty years. The arrest is huge news in and of itself, but for people interested in the social uses of genetic data, the way in which DeAngelo was identified—using genetic genealogy & genetic data from crime-scene samples—was noteworthy. Here, we discuss some of the genetics and math underlying the way in which he was identified (see also Henn *et al*. (2012)). Because there’s been lots of discussion of the ethics of these approaches, we will not focus on that here; see here for a collection of links & news articles.

The use of genetic data to identify suspects is not new. In the US, law enforcement makes extensive use of their CODIS (Combined DNA Index System) database—genetic searches against the database have aided almost 400,000 investigations since the mid-1990s. The CODIS database contains the genotypes of over 13 million people, most of whom have been convicted of a crime. The genetic information included about each person in the CODIS database is relatively sparse. Most of the profiles record genotypes at just 13 sites in the genome (since 2017, 20 sites have been genotyped). Because the CODIS sites are highly variable microsatellites, CODIS genotypes identify people nearly uniquely—they are sometimes called DNA fingerprints. (The CODIS markers reveal more than fingerprints do, though–they can reveal considerable ancestry information (Algee-Hewitt *et al*., 2016), can reveal close relatives (Rohlfs *et al*., 2013), and in some cases, its possible to identify genome-wide genetic profiles that match a particular CODIS dataset well (Edge *et al*., 2017).)

In a typical case in which law enforcement uses genetic data, the procedure is to genotype a crime-scene sample at the CODIS loci and look for a full or partial match against the CODIS database. If the sample came from a person who is in the CODIS database, he or she is likely to be identified. If there is no match, then the genetic search ends unless other information can be brought to bear.

In the Golden State Killer case, genotyping the samples at the CODIS markers did not reveal a match—Joseph DeAngelo was apparently not included in the CODIS database. Nonetheless, the genetic search continued. Investigators apparently genotyped the crime scene sample at a genome-wide set of SNPs, or single-nucleotide polymorphisms. SNPs are the markers of choice for large consumer genetics services like Ancestry and 23andMe (as well as for genome-wide association studies, GWAS). The police cannot access private databases like these—at least not without an extended legal process—but they do not have to. Many users upload their SNP data to third-party websites to perform advanced analyses or to search for matches with people tested by different companies.

These SNP databases are growing rapidly. Figure 1 shows the number of users in each of a set of repositories over the last few years (plot from here). The largest databases—AncestryDNA and 23andMe— are private. But the fourth-largest—GEDmatch, which now has about 950,000 profiles (Lussier and Keinan, 2018)—is an online service that searches for genetic matches with any user who uploads an appropriately formatted genotype file. Thats the one that police searched for DeAngelo.

**Figure 1:**
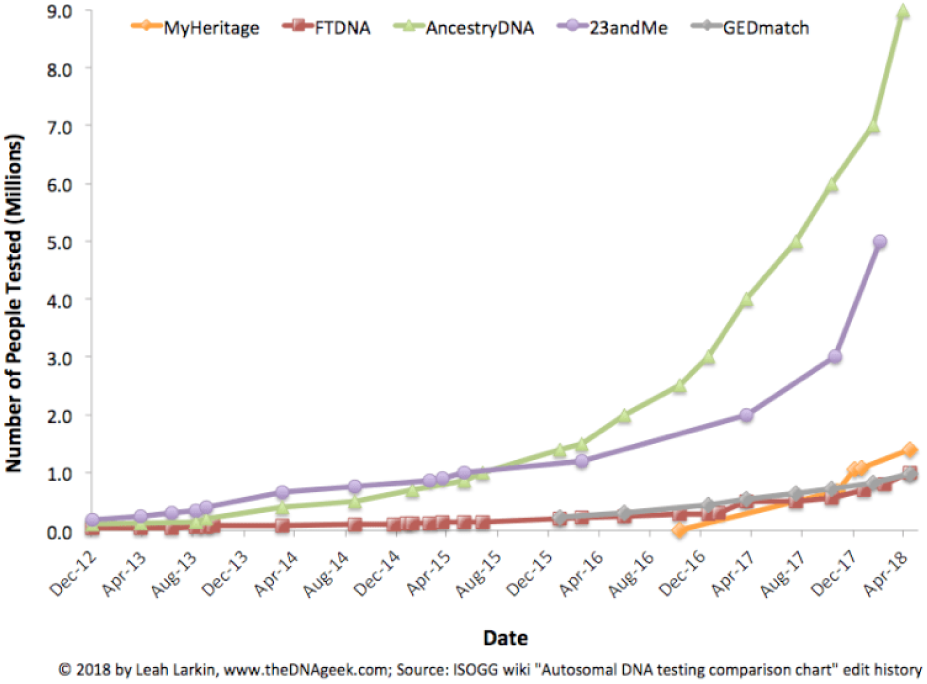
The numbers of people included in various genetic genealogy databases have grown rapidly in the past few years. Image courtesy of Leah Larkin.

Investigators searched for the suspects profile by making a personal user account and uploading a genotype file created from the SNP data obtained from crime-scene samples. To do this, the investigators must have created a data file mimicking the SNP set and file format provided by some genetic genealogy company. There was no exact match in the GEDmatch database—indeed, investigators did not expect that DeAngelo would have uploaded his own data—but the trail was not yet cold. The police could still run a search scanning the database for relatives of the suspect. If it is possible to identify a close relative, then the search for the suspect will be narrowed considerably, even if the suspect is not in the database. This is similar to the familial searching done using the CODIS database, which is legal in some states. (But it is imperfect, see Rohlfs *et al*. (2013) and Rohlfs *et al*. (2012).)

However, in the CODIS database, familial search efficacy is limited to close relatives (usually parents and siblings, and more tenuously uncles/aunts/nieces/nephews and first cousins). Thirteen microsatellite markers worth of information is simply not enough to distinguish a distant cousin from an unrelated person. With the hundreds of thousands of markers on a typical SNP chip, familial searching is much more powerful—third cousins can be found most of the time, and many (but not all) fourth cousins can be found too. A sample set of profile matches from GEDmatch is shown in Figure 2.

**Figure 2:**
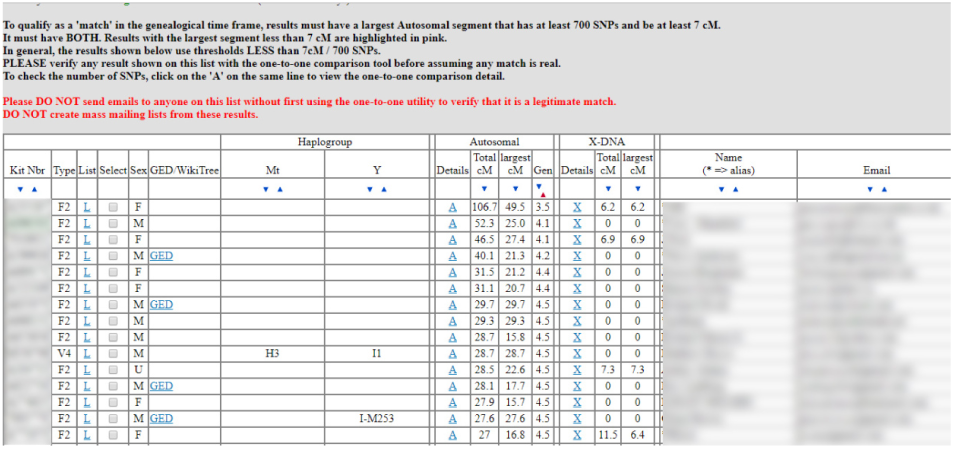
An anonymized screenshot of an example set of matches provided by GEDmatch.

Looking at SNP-based relative matches in GEDmatch, police found what they needed in the form of 10 to 20 likely relatives. These likely relatives represented third-to-fourth cousins of DeAngelo, most of whom he had probably never met. Using this genetic data, in combination with genealogical information about these relatives, the Golden State Killer investigation narrowed to one extended family, eventually honing in on DeAngelo himself.

Geneticists and genetic genealogists have been using these techniques for some time; the GEDmatch database exists because genealogists wanted to share genomic resources to help identify relatives, allowing families to be reunited (see here). Widespread reporting of the method used to identify DeAngelo as the suspected Golden State Killer has inspired a surge of interest in genetic privacy (see Erlich and Narayanan, 2014, for a general review of topic). Though DeAngelos capture is widely celebrated, people are also understandably surprised that the decisions of third or fourth cousins can potentially expose one to surveillance. In this post, we explore some simple models to ask questions about the extent of surveillance that is possible using the methods employed in the Golden State Killer case.

Two opposed phenomena govern the effectiveness of familial searches on genetic databases, one genealogical and one genetic. The genealogical phenomenon, which we could call “genealogical blowup,” is that the number of relatives one has at a specified degree of relatedness increases as the relatedness becomes more distant. For example, whereas a typical person may have one, two, or three siblings, he or she will usually have a large number—dozens or even hundreds—of third cousins (or third-degree cousins). Figure 3 shows the genealogical blowup phenomenon. On the left, we see the probability that a random person has at least one cousin of degree p in a database (depending on the size of the database), and on the right, we see the average number of cousins contained in a database. The number of genealogical cousins one has—where genealogical cousins are cousins in the usual sense, those connected by genealogy—increases rapidly for more distant relationships.

**Figure 3:**
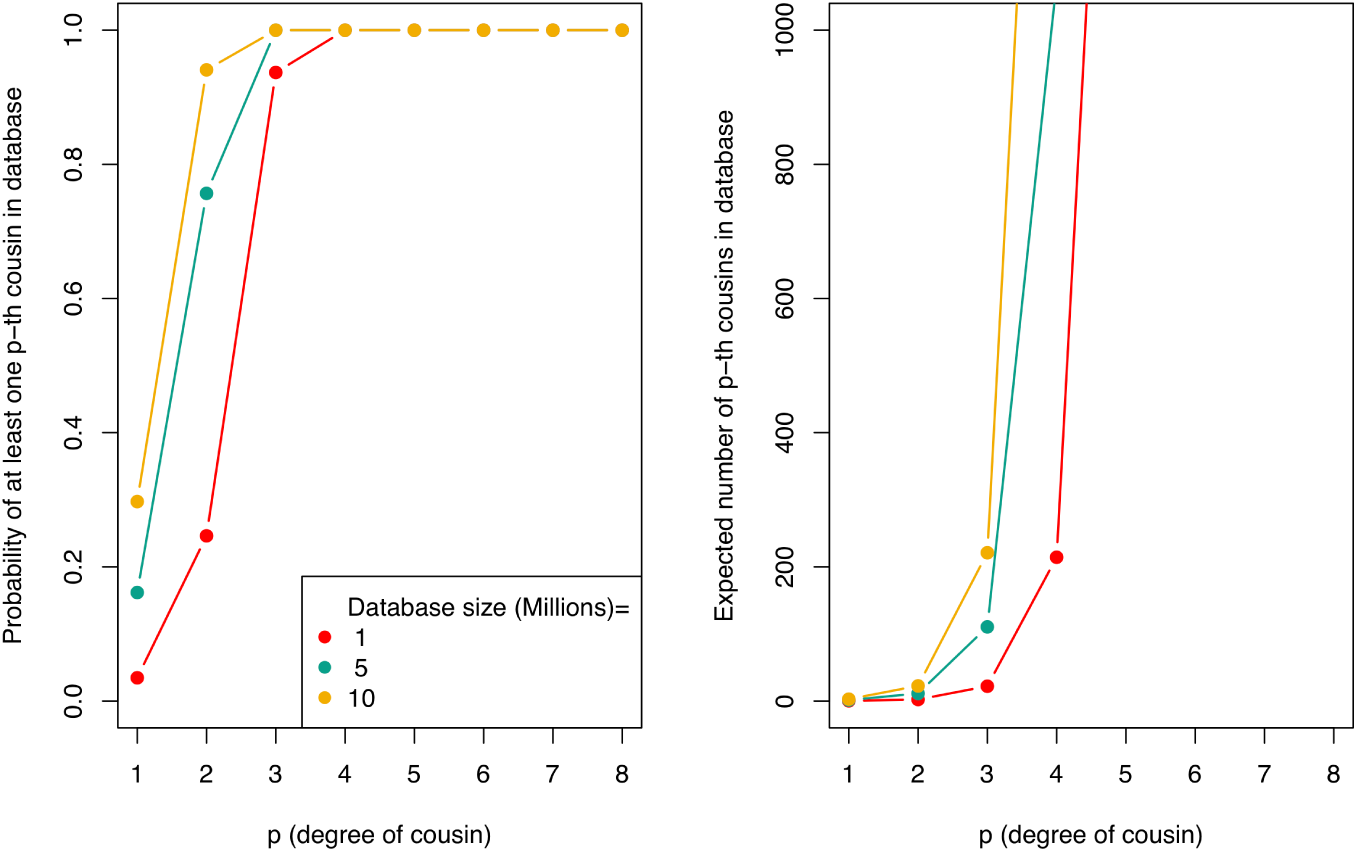
The probability of a random person having at least one cousin in the database, and the expected number of cousins in the database, as a function of the degree of cousin, *p*, and database size. The calculation on the left is based on the work of Shchur and Nielsen (2018). To make our calculations, we adopt some simplifying assumptions that are certainly wrongnamely complete inbreeding avoidance, monogamy with random mating, non-overlapping generations, random participation in the database, and population sizes similar to US census sizes across the last few generations. However, these calculations are useful to get a rough sense of the problem. Some details and pointers to other sources are in the notes below. The primary caveat that our assumptions entail is that our computations apply most directly to ancestry groups that are well represented in the database. GEDmatch is mostly composed of profiles from Americans of European ancestry. Recent immigrants to the US and people from non-European backgrounds are likely to find fewer relatives in GEDmatch than are European-Americans whose families have been in the US for a few generations.

The opposing genetic phenomenon is the noisiness of genetic inheritance (see Donnelly, 1983; Huff *et al*., 2011). Whereas the typical person has many distant cousins, the amount of genetic material shared with each of these distant cousins is small. You are nearly certain to share a lot of your genome with your first cousin, as you both have inherited a lot of your genomes from your shared grandparents. As a result, it is easy to identify pairs of first cousins if they are in the database.

The genomic material you share with your first cousin is the overlapping fragments of genome that both of you have inherited from your shared grandparents. In Figure 4 we show a simulation of you and your first cousins genomic material that you both inherited from your shared grandmother (details about how we made these simulations here). In the third panel we show the overlapping genomic regions in purple. These are regions where you and your cousin will have matching genomic material, due to having inherited it identical by descent from your shared grandmother. (If you are full first cousins, you will also have shared genomic regions from your shared grandfather, not shown here.)

**Figure 4:**
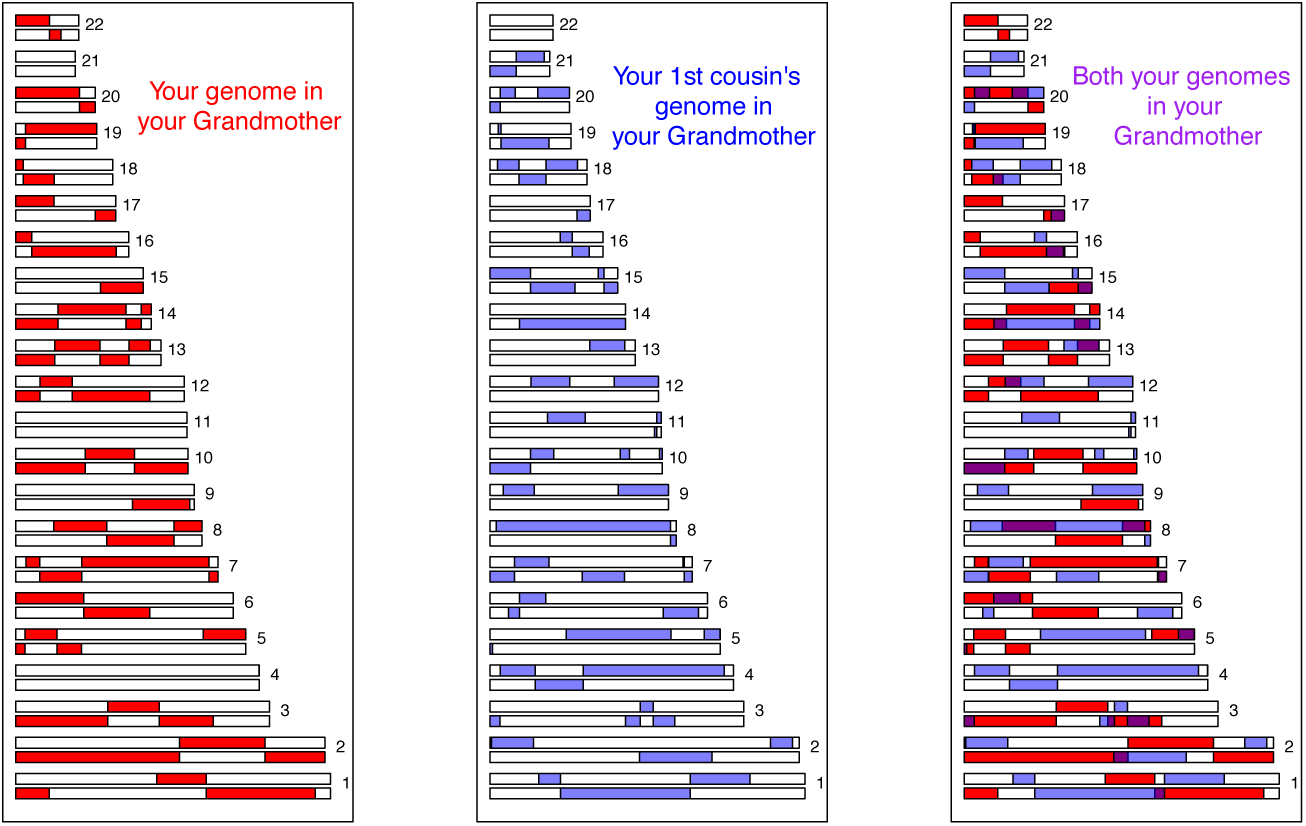
The degree of genetic overlap (identity by descent) for a pair of first cousins in one simulation. Shared regions from one of two shared grandparents are shown. The left and center panels show each cousin’s inheritance from the shared grandparent, whereas the right panel shows both, with regions where both cousins inherited material from the same grandparent—and thus will be identical by descent— colored in purple.

Now consider the case of third cousins. You share one of eight sets of great-great grandparents with each of your (likely many) third cousins. On average, you and your third cousin each inherit one-sixteenth of your genome from each of those two great-great grandparents. This turns out to imply that on average, a little less than one percent of your and your third cousins genomes (2 × (1/16)^2^ = 0.78%) will be identical by virtue of descent from those shared ancestors. If you do share one percent of your genomes, then your relationship to your cousin will likely be detectable using SNPs—the shared portions will be concentrated in relatively long stretches of chromosome that are easy to see statistically. But the more interesting thing is the variation around that average. There is a non-trivial chance (about 2%) that you will actually share no identical segments of your genome with your third cousin—in that case, we say you are genealogical cousins but not genetic cousins.

Figure 5 shows an example where third cousins share some blocks of their genome (on chromosome 16 and 2) due to their great, great grandmother. While Figure 6 show an example where the same individual shares the same great, great grandmother with another 3rd cousin, but has no genetic sharing due to that connection.

**Figure 5:**
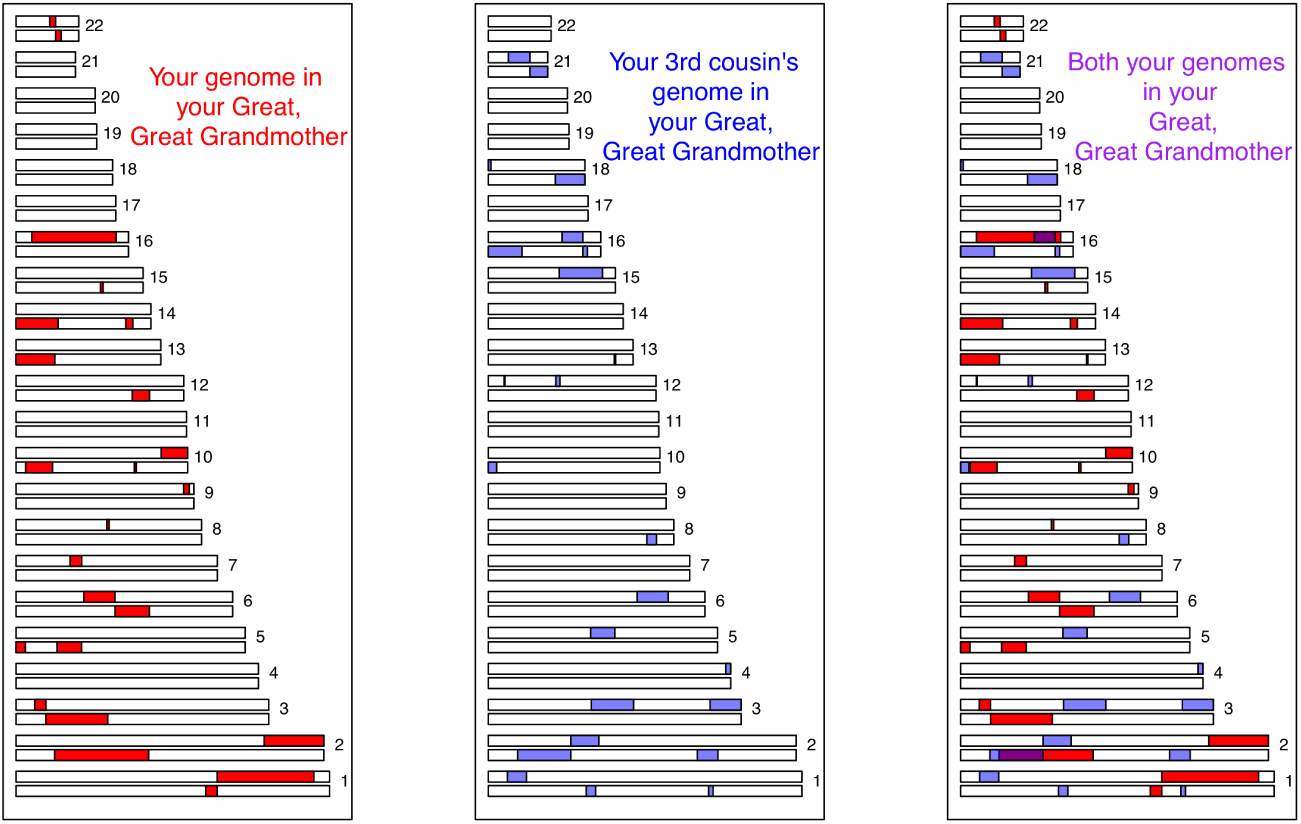
The degree of genetic overlap (identity by descent) for a pair of third cousins in one simulation. Shared regions from one of two shared great-great grandparents are shown. The left and center panels show each cousin’s inheritance from the shared great-great grandparent, whereas the right panel shows both, with regions where both cousins inherited material from the same ancestor—and thus will be identical by descent—colored in purple.

**Figure 6:**
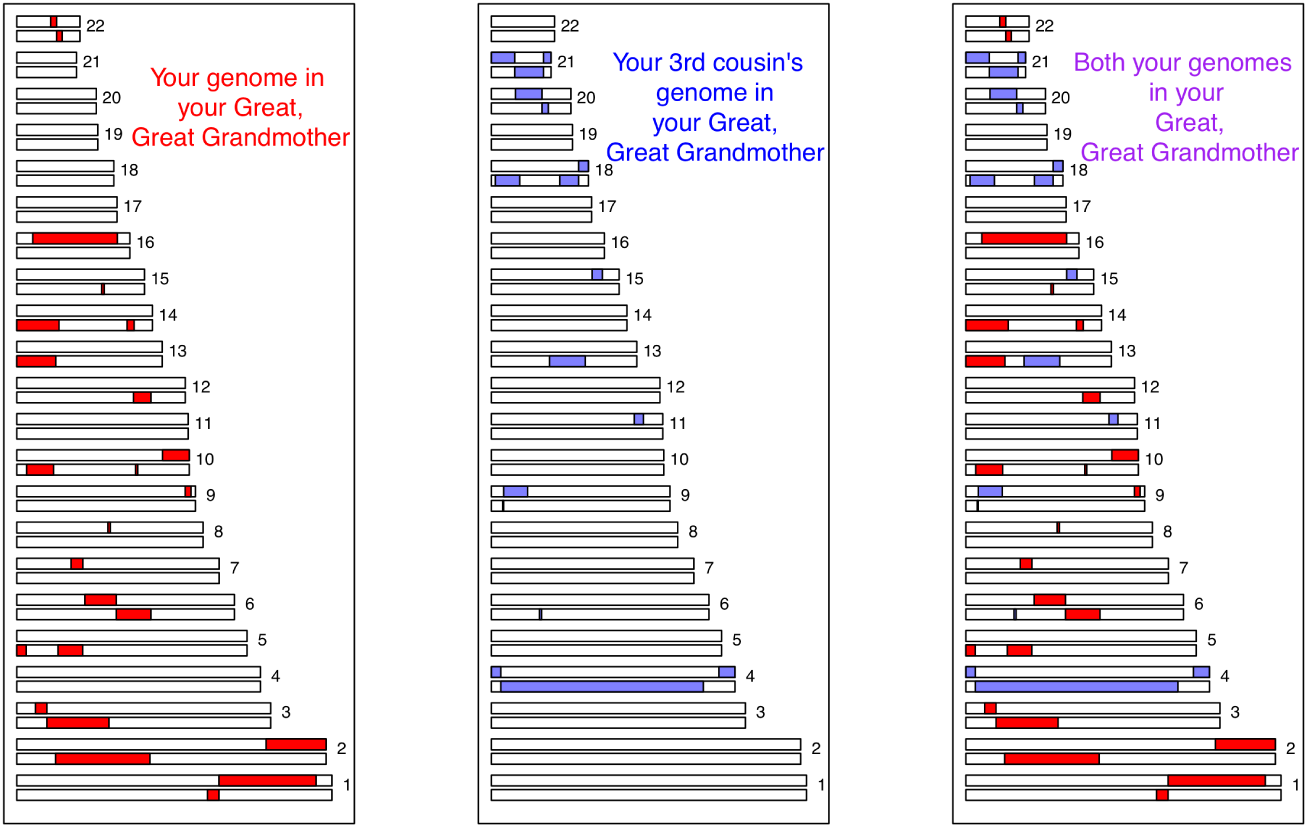
Another simulation of third cousins—in this simulation, the third cousins share no genetic material identical by descent by virtue of their shared ancestor.

As the degree of relatedness decreases—on to fourth cousins, fifth cousins, and so on—an ever-larger proportion of ones genealogical cousins will not be genetic cousins. Figure 7 shows the proportion of degree-*p* cousins with which one expects to share either at least one, two, or three genetic blocks. Sharing 1 block is not very informative (see here). Individuals with whom one shares three or more large genetic fragments are likely strong leads. (Again, the assumptions used here are explained in the notes below.)

**Figure 7:**
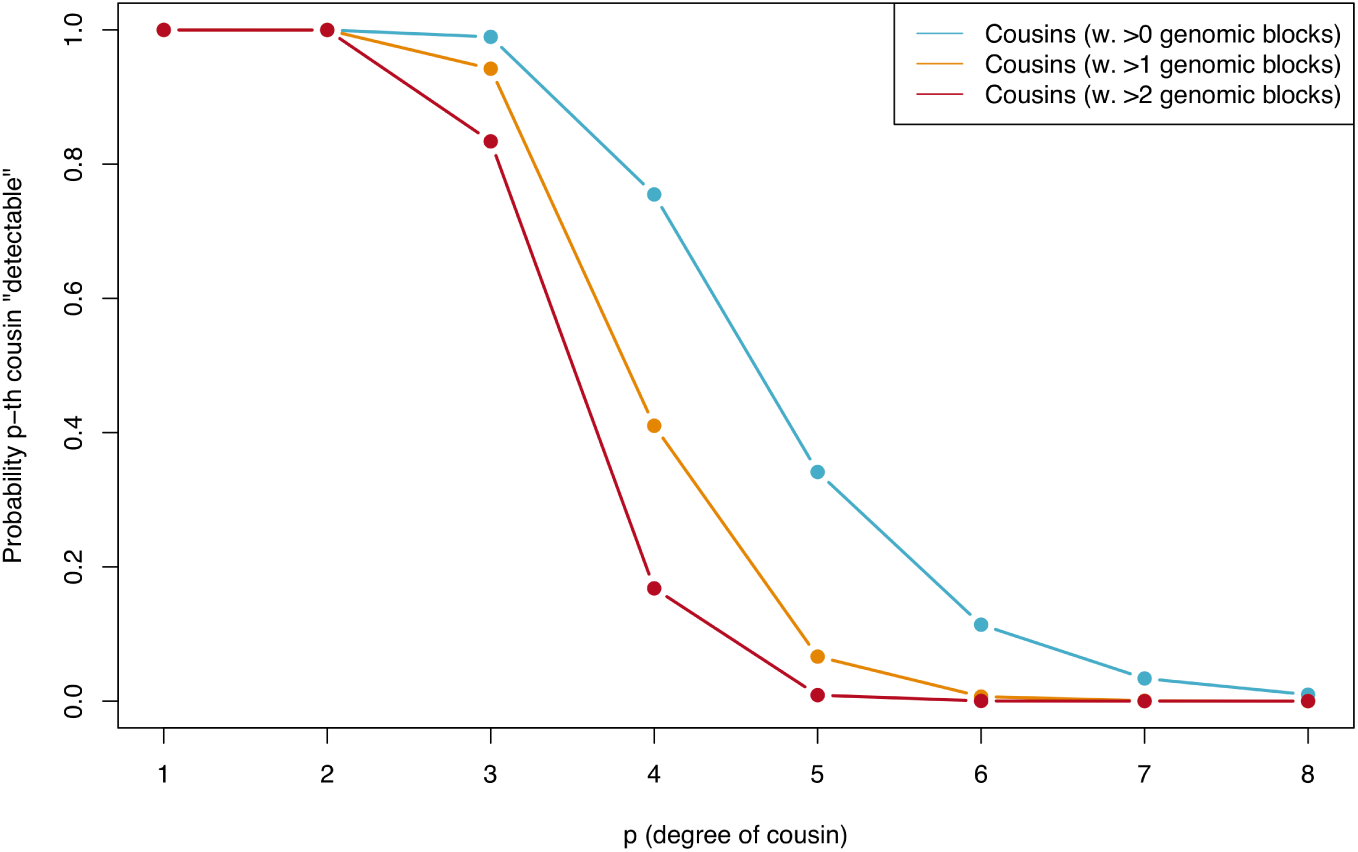
The probability that a cousin of degree *p* is detectable genetically as a function of *p*. Genetic detection depends on the threshold one sets for declaring a pair of people related (different colored lines).

An appreciation of these two phenomena—genealogical blowup and the noisiness of genetic inheritance—is crucial for understanding how public SNP databases might be used by law enforcement in the future. There is a tradeoff. One typically has a large number of genealogical eighth cousins, but only a small proportion of them will be genetic cousins, and even these are often impossible to identify as such. On the other hand, it is easy to detect ones first cousins, but because one typically has a small number of first cousins, the probability that a random person has one in a genetic database is low unless the database is very large. (Another factor relevant for law enforcement is that closer matches are more useful; they narrow the pool of possible suspects more.) Figure 8 combines the considerations illustrated in the previous plots, showing the expected numbers of genetic cousins in the database. The tradeoff of genealogical blowup and the noisiness of genetic inheritance is optimized in the third to fifth cousin range—you have a lot of genealogical cousins at this degree of relatedness, and many of them will be detectable genetic cousins. Because closer relatives are more useful to law enforcement than more distant relatives, it’s likely that many of the cases that could be solved by these methods would involve some mix of 2nd, 3rd, and 4th cousins.

**Figure 8:**
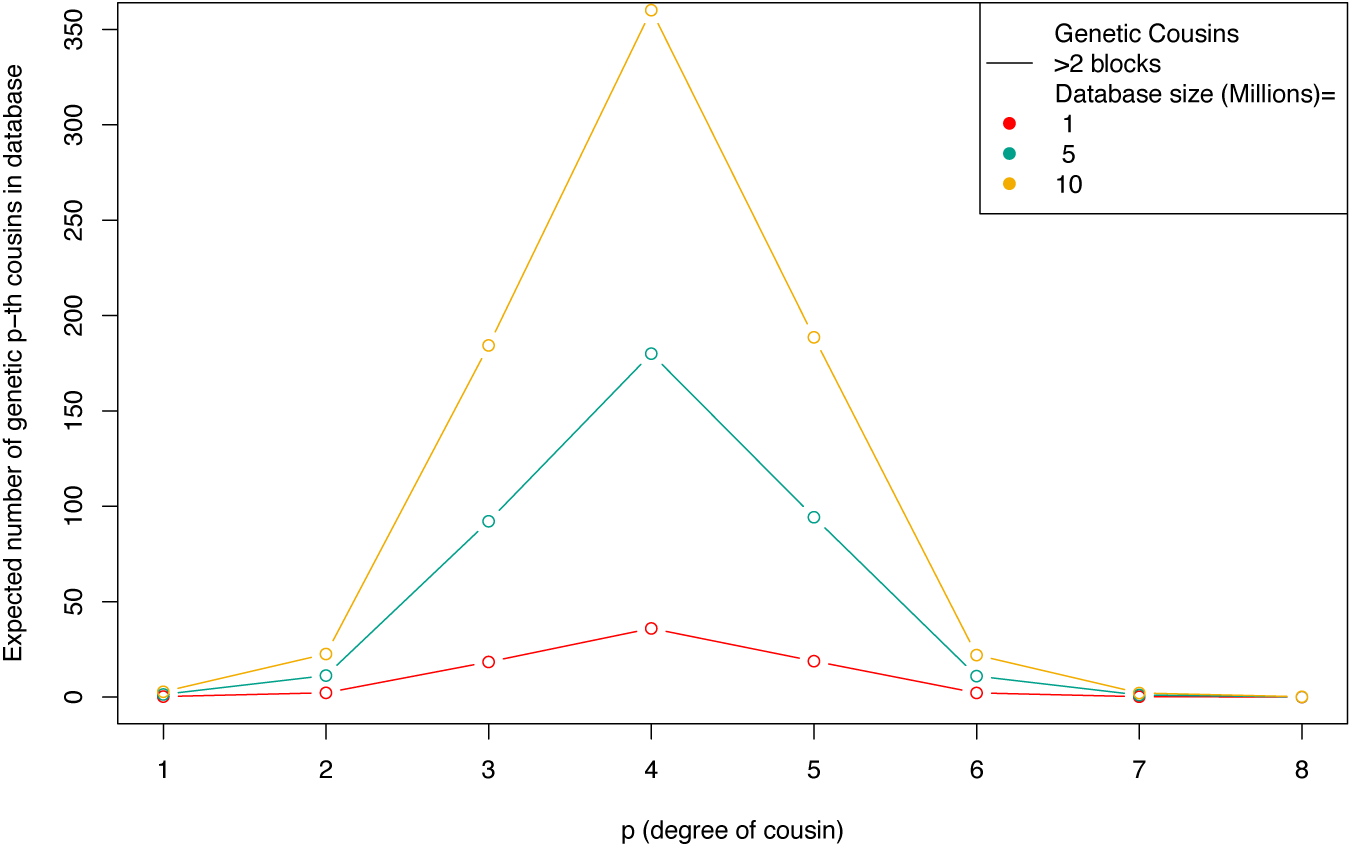
The expected number of genetically detectable cousins in a database (where a cousin match is declared if greater than two genetic blocks are identical by descent) as a function of the degree of cousin (horizontal axis) and database size (different colored lines).

The Golden State Killer results are close to what we expect given the size of the GEDmatch database. Under the assumptions we make here, its likely that a large percentage of people have at least one high-confidence genetic cousin in GEDmatch, and the number of 3rd-4th cousins found for DeAngelo—10 to 20—is not too far from the expectations. Its striking that uploading ones information to a matching database potentially opens up a large number of other people to eventual identification, and that most of these people are distant enough relatives that one would likely never have met them. To illustrate, consider that 13 million individuals in CODIS likely wouldnt reveal a familial match because only very close relatives are detectable in CODIS. But using the far smaller GEDmatch database (approximately 1 million people), investigators tracked DeAngelo down. As Yaniv Erlich put it recently, You are a beacon who illuminates 300 people around you. Its also striking that were already in an era in which familial searches against publicly accessible SNP databases are feasible for a lot of cases, probably the majority of cases where the suspect has substantial recent ancestry in the US—the public datasets are big enough (or will be soon). The limiting factor here may be the genealogical work to trace distant cousins through family trees, but big public datasets might make the genealogical task easier too. From here, its a question of deciding the circumstances under which we as a society want these familial searches to be used.

## Acknowledgments

Thanks to the Coop lab and Debbie Kennett for helpful comments on an earlier draft.

## Notes

A *p*th cousin is a person with whom one shares an ancestor (in our model, an ancestral couple) *p* + 1 generations ago (your great(*p* – 1) grandparents). If theres no inbreeding in ones recent family tree, then one is descended from 2*^p^* ancestral couples *p* + 1 generations ago. A pair of individuals in the present are *p*th cousins (or closer) if their sets of 2*^p^* ancestral couples overlap—they share ancestors *p* + 1 generations ago. Lets assume that there are *N*_*p*_ potential ancestors in *N*/2 couples, *p* generations back. If each of these couples have the same probability of having children and there is not too much variation in family size, we can view the problem as if people in the present choose their ancestors *p* + 1 generations ago at random. Your ancestors were no doubt very special people, but as far as this model is concerned they were just 2*^p^* random draws from all the couples whove left descendants. To calculate the probability that you and I are *p*th cousins, we just need to calculate the probability that our two sets of 2*^p^* ancestors overlap (note that this assumes monogamy, i.e. that well be full not half cousins, but even if that wasnt true, that just alters things by a factor of two). Now, we have something close to a classic probability problem: we draw a set of 2*^p^* balls at random from an urn with *N*_*p*_ balls, replace the balls in the urn, and repeat the draw of 2*^p^* balls—what is the probability that at least one ball is a member of both sets of 2*^p^* balls?

The probability that you and I are *p*th cousins is roughly (4*^p^*/(*N*_*p*_/2)), when *N*_*p*_≪ 2*^p^* i.e. when your ancestors are a small fraction of the total people in the population. In a current-day database of *K* individuals, drawn from the same population as you, your expected number of *p*th cousins is *K* × 4*^p^*/(*N*_*p*_/2). Two factors make this blow up quickly back over the generation. First, 4*^p^* grows quickly back over the generations; second, population sizes have increased rapidly in the recent past, which means that *N*_*p*_ declines quickly with *p* (because *p* counts generations backward in time).

One of biggest uncertainties in our calculations is the size of the pool of possible ancestors. Our calculations should therefore be viewed as crude approximation. Our calculations are based on assuming that the population size of possible ancestors is given by the census population size of the USA. To get the census population size we assume a generation time of 30 years, and take the population size in the decade 1950 – 30×(*p*+1). We assume that roughly ½ of the individuals in the population are potentially parents, and that 90% of potentially parents have children. We impose a floor on the population size that it cannot drop below 1 million potential parents, to reflect the fact that for people of European ancestry, the pool of ancestors back then would also include Europe. Given the large variation in family sizes N should likely be lower still, as variation in family size decreases the effective *N* further.

SHCHUR and NIELSEN (2018) recently worked through the probability that you have no *p*th cousins in a database of *K* individuals, in a model similar to that described above. The model Shchur and Nielsen (2018) use is more realistic than the one we consider here—it allows for some inbreeding and takes explicit account of the fact that some couples will not have children. They find (their equation 7) that the probability that an individual has no *p*th cousins in the database, given a fixed population size of *N*, is approximately exp(–2^(2×*p*−2)^ × *K*/*N*).

The math underlying the genetic calculation is described in more detail here (see Donnelly, 1983; Huff *et al*., 2011). To summarize: if you share two ancestors *p* + 1 generations with your *p*th cousin, then you share a particular autosomal chromosomal region with probability 2 × (1/2^*p*+1^ − 1). You have 22 autosomal chromosomes, and each generation, recombination happens in ∼34 places on these chromosomes. Looking back *p* + 1 generations, your chromosomes are broken up into approximately (22 + 34 × (*p* + 1)) chunks, which are spread across your ancestors. Likewise, your relatives genome is broken into (22 + 34 × (*p* + 1)) chunks. Because recombination events rarely happen in the exactly same place, your two genomes combined are broken into (22 + 34 × *d* × 2) pieces. As each of these is inherited identical by descent to both you and your cousins from that ancestor with probability 1/2^2(*p*+1−1)^, you and your cousins should expect to share *E_B_* = 1/2^2(*p*+1)−1^2 × (22 + 34(*p* + 1)) blocks of your autosomal genome. The probability that you share 0 blocks is approximately exp(−*E_B_*), while the probability of sharing 2 or more blocks (*Q*_*p*_) can approximately be obtained under the Poisson distribution (which is a good approximation beyond 1st cousins).

Putting all of this together, your expected number of genetic pth cousins is (*Q_p_* × K × 4^*p*^/(*N_p_*/2). Thats the solid line plotted in the final figure.

